## Supplementary material for "Sexual selection theory meets disease vector control: Testing harmonic convergence as a “good genes” signal in *Aedes aegypti* mosquitoes": SI

### Materials and methods

#### *Parental rearing*

Mosquito eggs were vacuum hatched for 20 min and received a pinch of powdered fish food diet (Hikari Cichlid Gold, Hayward, CA, USA). Hatched larvae were held at 27 °C and 85% relative humidity (RH) overnight. The following day larvae were sorted into groups of 200 and placed in 1 L of distilled water in plastic trays (28 x 21 x 8 cm) to obtain medium body sizes [1]. Each tray was provided with four large pellets of fish food and monitored daily. Fitness differences often remain undetected under optimal rearing conditions, and environmental stress has been shown to be important for measuring genetic benefits in many animals [2–4]. Because we previously found that differences between offspring of converged and non-converged males are detectable only after application of a moderate level of temperature stress [5], we therefore exposed $4^{th}$ instar larvae to 21 ºC ambient temperatures for 24 h. Resulting pupae were individually placed in 15 mL tubes plugged with cotton wool. Upon emergence, males and females were held in sex-specific cages (8L plastic buckets) where they fed a 10% sugar solution *ad libitum*.

In all experiments, mosquito wing lengths were obtained from a subset of mosquitoes from each experimental group as a proxy for overall body size as previously described [6]. For reproductive fitness experiments, regardless of whether parent or offspring data were used in subsequent analyses, father (N=90) and mother (N=84) wing lengths were 2.23 ± 0.10 and 2.96 ± 0.12 mm, respectively, and son (N=72) and daughter (N=139) wing lengths were 2.42 ± 0.34 and 2.93 ± 0.10 mm, respectively. For flight performance experiments, father (N=34) wing lengths were 2.13 ± 0.13 mm.

#### *Harmonic convergence assay*

Females were tethered using the "semi-tethering" technique to optimize flight mobility while maintaining female proximity to the microphone as previously described [5,7,8]. Briefly, females were tethered at the dorsal mesothorax by a human hair attached to an insect pin using nail glue (L.A. Colors, Ontario, CA, USA or similar product). Tethered females were placed approximately 3 cm away from a particle velocity microphone (NR-21358; Knowles Electronics, Itasca, IL, USA) attached to a custom amplifier [9] in a plastic recording arena (20 cm x 14 cm x

10 cm) [10]. Female flight was stimulated by using an aspirator to release females from tarsal inhibition or to deliver brief pulses of air [5,8]. All parental recordings were performed between the hours of 09:00–18:00 and the temperature and humidity conditions were 26.47 ± 1.13˚C and 59.01 ± 7.95% RH for the reproductive fitness assays and 25 ± 1.8°C and 68.1 ± 2.0% RH for the flight performance assays.

Potential instances of harmonic convergence were systematically tested at all harmonic combinations below 3,000 Hz, including at the female second and male first harmonics (2:1), the female third and male second harmonics (3:2), the female fourth and male third harmonics (4:3), the female fifth and male third harmonics (5:3), and the female fifth and male fourth harmonics (5:4) [11]. To be considered a true instance of convergence, the harmonics had to be within 5 Hz of each other for a duration of at least 1 s [5]. If no courtship flight interactions occurred within 5 min, we considered it a non-event and started a new trial with a fresh set of mosquitoes.

***Offspring generation and rearing***

*Offspring reproductive fitness assays*

Eggs from females that took blood meals and survived to lay were collected from each mating pair, vacuum hatched, and reared separately. Larvae were reared in plastic cups containing approximately 25 eggs per family and 50 mL of deionized water. No more than 20 male pupae were placed directly into 0.5 L wax containers supplied with 10% sugar-soaked pads. Female daughter pupae were placed in individual 15 mL tubes. Females used in male insemination capacity experiments and males used to inseminate daughters for the lifetime reproduction study were reared from a separate cohort of unrelated Thai colony mosquitoes as described above for parents.

*Offspring flight performance assays*

For son flight performance assays, 20 larvae per family were held in 100 mL of water and larvae were fed diet *ad libitum.* Offspring pupae were placed into individual 15 mL tubes for adult eclosion. Upon eclosion, offspring were separated by sex and family. Female offspring were discarded while males were sorted into 0.5 L cups by emergence day and family and supplied with 10% sugar.

***Reproductive fitness assays***

*Male fertility: Insemination capacity*

In the case of fathers, the insemination capacity assay followed the initial 24 h parental mating period, in which the father was held with the original female. Each male was then held in an 0.5 L container with five virgin females for 48 h intervals. The spermathecae of females replaced at each interval were dissected and examined for the presence or absence of sperm at the end of each 48 h interval.

*Female fecundity: Eggs laid*

Upon emergence, individual daughters were transferred into 20 mL cartons and held for three days to allow for reproductive maturation. Daughters were then provided three unrelated colony males for 24 h to allow for insemination. After removal of males, females were offered a blood meal (co-author SAP) every three days for the remainder of their lifetime up to 20 days. Oviposition cups with water were added to each carton three days post-blood meal. Females that did not ingest the initial blood meal were discarded and data were recorded on whether females blood fed during each subsequent host offering. Consistent with feeding behavior of *Ae. aegypti* in nature [12,13], females were not provided sugar for the duration of the fecundity assay.

***Flight performance assay***

*Male flight performance: Mating attempts, contacts, and contact success rate*

For the custom-built flight response cage, the speaker was suspended 8 cm below the lid of a 30 $cm^3$ clear plastic cage and was attached to a timing belt (Synchroflex Timing Belt, 4660-950, Technobots, Rugby, UK) on top of the cage through a 0.5 cm x 21 cm track in the lid. The timing belt was driven in a back and forth motion across the cage using a small stepper motor (ST411, 18M1804A, Nanotec, Munich, Germany) powered by an enclosed power supply (LRS-150F-12RS, Mean Well Enterprises, Taipei, Taiwan). The entire apparatus of was placed on top of a hot plate set to 28˚C with a worn black t-shirt (co-author LJC) stretched across the top to provide additional visual and olfactory mating cues for males, as males tend to swarm around human hosts when seeking mates [14] and prefer dark colors [15].

Individual males were released into the cage with a silent, immobile speaker to allow for a 15–20 s cage acclimation period in the absence of acoustic stimuli or speaker movement. After this, the silent speaker began to move for an approximately 10 s during a movement acclimation period, with the arm taking 15 ms to accelerate to its 3 m/s back and forth speed across the cage. Finally, audio playback began and continued for two minutes during the moving playback period, where a 550 Hz pure tone playback was presented at a volume of approximately 60 dB in 10 s bursts with 5 s of silence in between. After the trial, males were removed from the flight response cage and held individually in 30 mL cartons at 27˚C and 80% RH overnight before testing in harmonic convergences assays.

Acoustic stimuli most likely to attract free flying males were identified the through a series of pilot playback experiments (S2 Fig.). The microphones and ear bud speaker were placed in the center of a 20 $cm^3$ cage and secured on top of a polystyrene platform (13 x 10 x 5 cm). A particle velocity microphone was placed 0.5 cm below the speaker and a pressure microphone (FG-23329-C05; Knowles Electronics, Itasca, IL, USA) was placed adjacent to this. For each trial, groups of 15 males were randomly collected from the colony cage and released into the flight cage. There was a 30 s period for the males to acclimatize before the playback began. The two-minute playback consisted of 15 s of pre-stimuli silence with three bursts of 15 s stimuli with 20 s of rest in-between each burst. For any one cage, each of the six frequency treatments were played twice in a randomized order, giving 12 recordings per cage. We recorded the number of males approaching a given frequency in each trial cage. This experiment was performed twice giving a total of 28 trials with 420 males. We identified 550 Hz as the frequency that most free-flying males responded to (S1 Fig).

A second pilot experiment was conducted to determine at what speed the movement of the speaker became discerning. Five males per trial were released into the flight response cage. The speaker was driven at 200, 300, 400, and 500 RPM. A 550 Hz tone was played for 10 s pulses with a 5 s rest period in between each pulse for a total of two minutes. Video of each trial was recorded using an HD Camera positioned parallel to the speaker movement. We recorded the number of attempts made by the five males and the number of males able to contact the microphone during this trial period. We replicated this assay three times per speed (5 males/speed/replicate). Males were most responsive to a speaker moving at 300 RPM. On average, there were 8.67 ± 3.38 mating attempts with 30.54% of these including a contact with the speaker. By contrast, the contact success rate for a stationary speaker playing the same stimuli is 100% and a speaker moving at 400 RPM is only 3.3%. While some mating attempts were made at all speeds, no males in the 200 and 500 RPM groups were successful at contacting the speaker.

#### Data analysis

*Distribution and variability of reproductive fitness and flight performance traits*

To characterize the distribution and variability of parent and offspring reproductive fitness and flight performance parameters, data were first tested for normality using a one-sample Kolmogorov-Smirnov (KS) test in SPSS. Distributions were then compared for equality using an independent samples Kruskal–Wallis (KW) test. Descriptive statistics of fitness and performance data were produced to determine skewness, standard deviation, and other dispersion statistics (Table 1).

*Heritability of reproductive fitness and flight performances traits*

To assess the heritability of the reproductive fitness and flight performances metrics, we ran linear mixed effects models (LMMs) [16,17] using a gaussian distribution to test for relationships between parental and offspring traits. In each case, the parental trait (father insemination capacity, father wing length, number of eggs in mother's first clutch, or mother wing length) and replicate were incorporated as fixed effects and family was incorporated as a random effect to account for the measurement of multiple offspring from a single parental pair. Fixed effects were tested using F tests with a Satterthwaite approximation for the denominator degrees of freedom and assumptions of normality and homogeneous variances were checked by visually assessing residuals. For reproductive fitness assays in sons, we assessed the effect of parental traits on son insemination capacity and wing length response variables. For reproductive fitness in daughters, we assessed the effect of parental traits on daughter eggs laid in first clutch, total eggs laid, eggs laid per blood meal, days alive, and wing length response variables. For flight performance assays, we used the same approach to test the relationship between father and son flight metrics, including total mating attempts, total contacts, and contact success rates using LMMs, and probability of contact using a generalized linear mixed model (GZLMM) [16].

*Harmonic convergences signaling of parent and offspring inherent quality*

To test whether harmonic convergence is used as an acoustic signal of reproductive fitness and inherent genetic quality, we first determined whether convergence status predicted fitness traits within the parental generation. Linear models (LM) [16] with harmonic convergence status, replicate, and wing length incorporated as fixed effects were used to test for differences in father insemination capacity and mother eggs laid in first clutch response variables. For LMs, fixed effects were tested using F-tests and assumptions of normality and homogeneous variances were tested as with LMMs. A LM with harmonic convergence status, replicate, and mating status as fixed effects was used to test the effect of convergence on father total attempts, total contacts, and contact success rate response variables. A generalized linear model (GZLM) [16] with a binomial distribution and the same effects was used to assess the impact of convergence status on the probability that the father successfully contacted the speaker at least once in the trial. For GZLMs, fixed effects were tested using t-tests and z-tests.

Next, to examine whether harmonic convergence cues serve as indicators of indirect effects on offspring fitness, we tested whether parental convergence status predicted offspring fitness measures. For sons, LMMs [18] with parental convergence status, replicate, and wing length as fixed effects and family as a random effect were used to test the effect of parental convergence on son insemination capacity (response variable). We also tested whether son convergence status predicted son insemination capacities using the same models. We used LMMs [16] to assess whether the fixed effects of parental convergence status and replicate or the random effect of family had an effect on son total mating attempts, total contacts, contact success rate, or probability of contact response variables. Here, as before, a gaussian distribution was used in all LMMs, except probability of speaker contact data (GZLMM), which used a binomial distribution. For GZLMMs, fixed effects were tested using z-tests or likelihood ratio chi-square tests (aka deviance tests). To test for an association between parental and son convergence status (response variable), we also used a GZLMM with a binomial distribution, replicate as a fixed effect, and family as a random effect.

For daughters, LMMs [18] with parental convergence status and replicate as fixed effects and family as a random effect were used to assess the effect of parental convergence on daughter eggs laid in first clutch, total eggs laid, eggs laid per blood meal, and days alive response variables. Kaplan-Meier curves were used to visually compare daughter survival based on parental convergence status [19–21]. A Cox frailty model was used to test for differences in survival between the curves while controlling for replicate as a fixed effect, cumulative blood meals as a time-varying fixed effect, and family as a random effect [22,23]. We additionally used the individual day level data to construct a daughter life table [24–27]. Using these data, we calculated the intrinsic rate of increase ($r$), cumulative reproductive rate ($R_0$), and generation time ($T_c$) for offspring of converged and non-converged parents from the two life table experiment replicates.

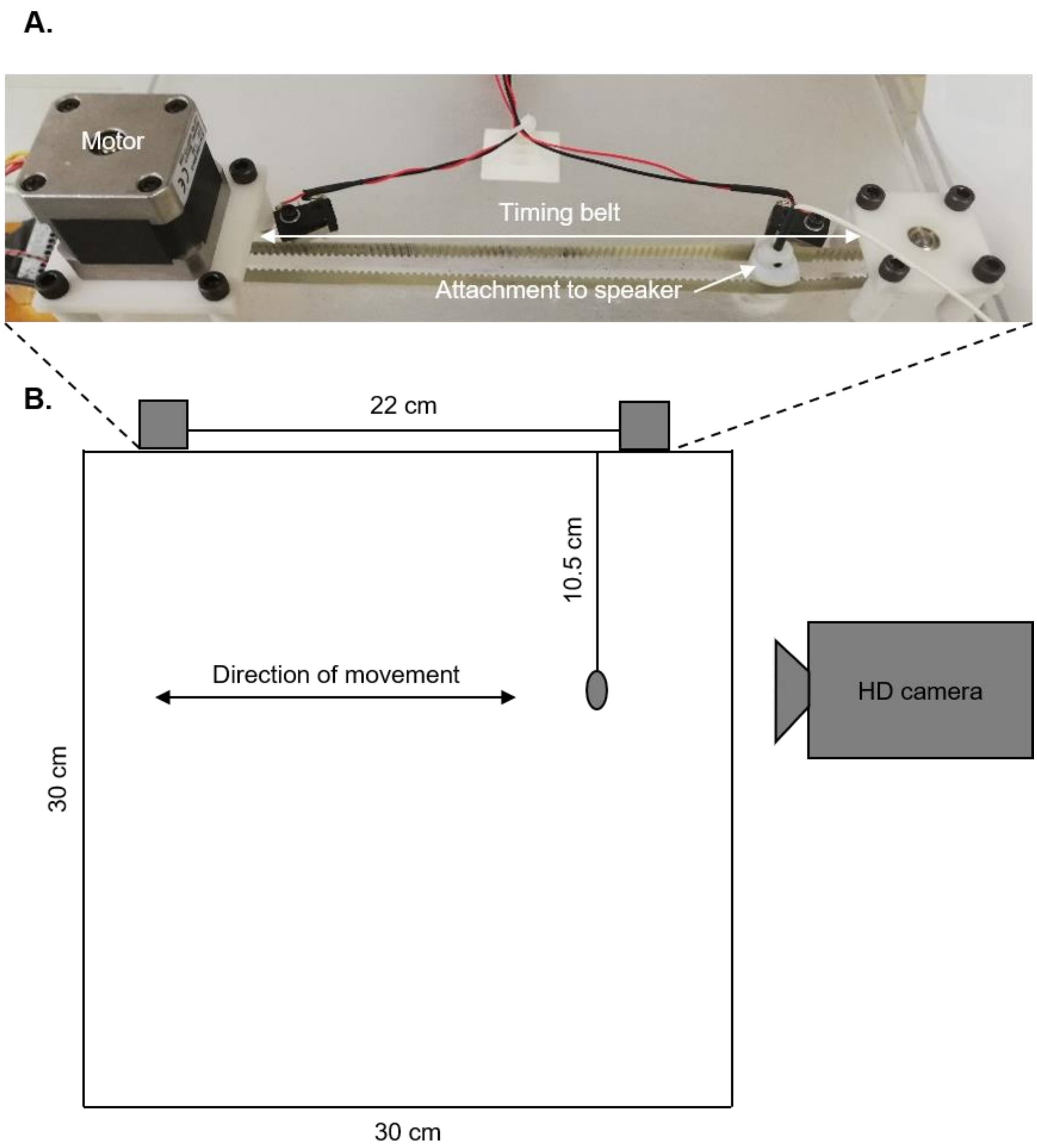


**S1 Figure. Flight performance assay set up.** A custom-built flight cage was fit with a moving arm that allowed for playback of stimuli from a moving source. A motorized timing belt attached to a speaker (A) was used to move the arm back and forth at fixed speeds. An HD camera was positioned parallel to speaker movement to capture male responses in the flight cage (B).

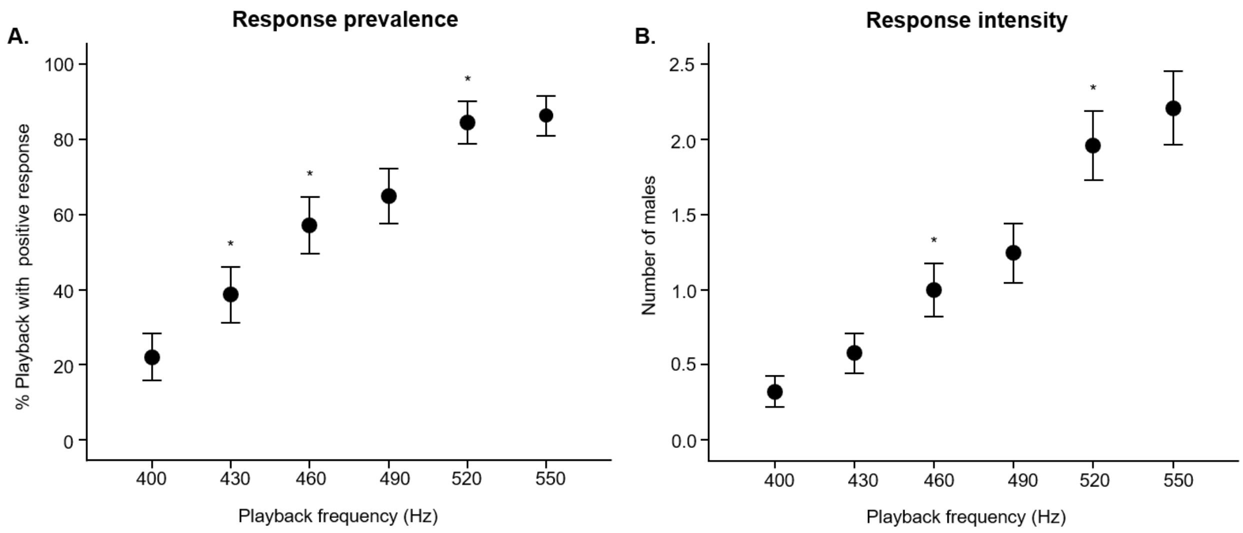


**S2 Figure. Flight tone parameterization for flight performance assay playback.** The effect of playback frequency on the response prevalence (A), or the proportion of playbacks at a particular frequency to which any male free-flying male was attracted to the speaker (LMM, P<0.001), and the response intensity (B), or the number of males (out of 15) that responded to playback at a particular frequency. The 550 Hz playback frequency was selected for flight performance assays since it was the tone that induced the most male responses. All error bars represent +2 standard errors.

**S1 Table. Daughter life table parameters do not differ by parental harmonic convergence status.** Daughter life table analysis data presented by parent convergence status and replicate (individually and averaged). Although the effect of parental convergence status on $R_0$ and $r$ (but not $T_c$) are inconsistent across two replicates, the average effect was similar. Abbreviations: N, sample size; $R_0$, cumulative reproductive rate; $T_c$, generation time; $r$, intrinsic rate of increase.

| Replicate | Parental convergence status | N | $R_0$ | $Tc$ | $r$ |
|---|---|---|---|---|---|
| 1 | Converged | 24 | 118.604 | 14.154 | 0.494 |
| | Non-converged | 74 | 90.815 | 15.117 | 0.450 |
| 2 | Converged | 52 | 108.385 | 14.560 | 0.452 |
| | Non-converged | 57 | 123.921 | 14.628 | 0.468 |
| Average | Converged | 76 | 113.494 | 14.357 | 0.473 |
| | Non-converged | 131 | 107.368 | 14.873 | 0.459 |

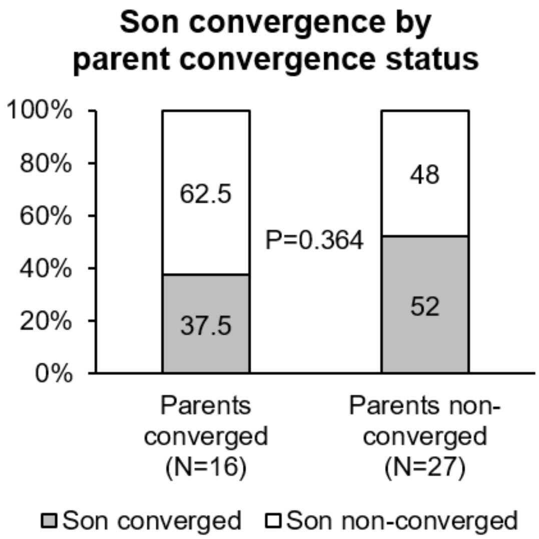


**S3 Figure. Parental harmonic convergence does not signal offspring convergence.** Son convergence status did not differ between those descending from converged parents and those descending from non-converged parents (P=0.364). Graphs display sample sizes (N) and GZLMM P-value.

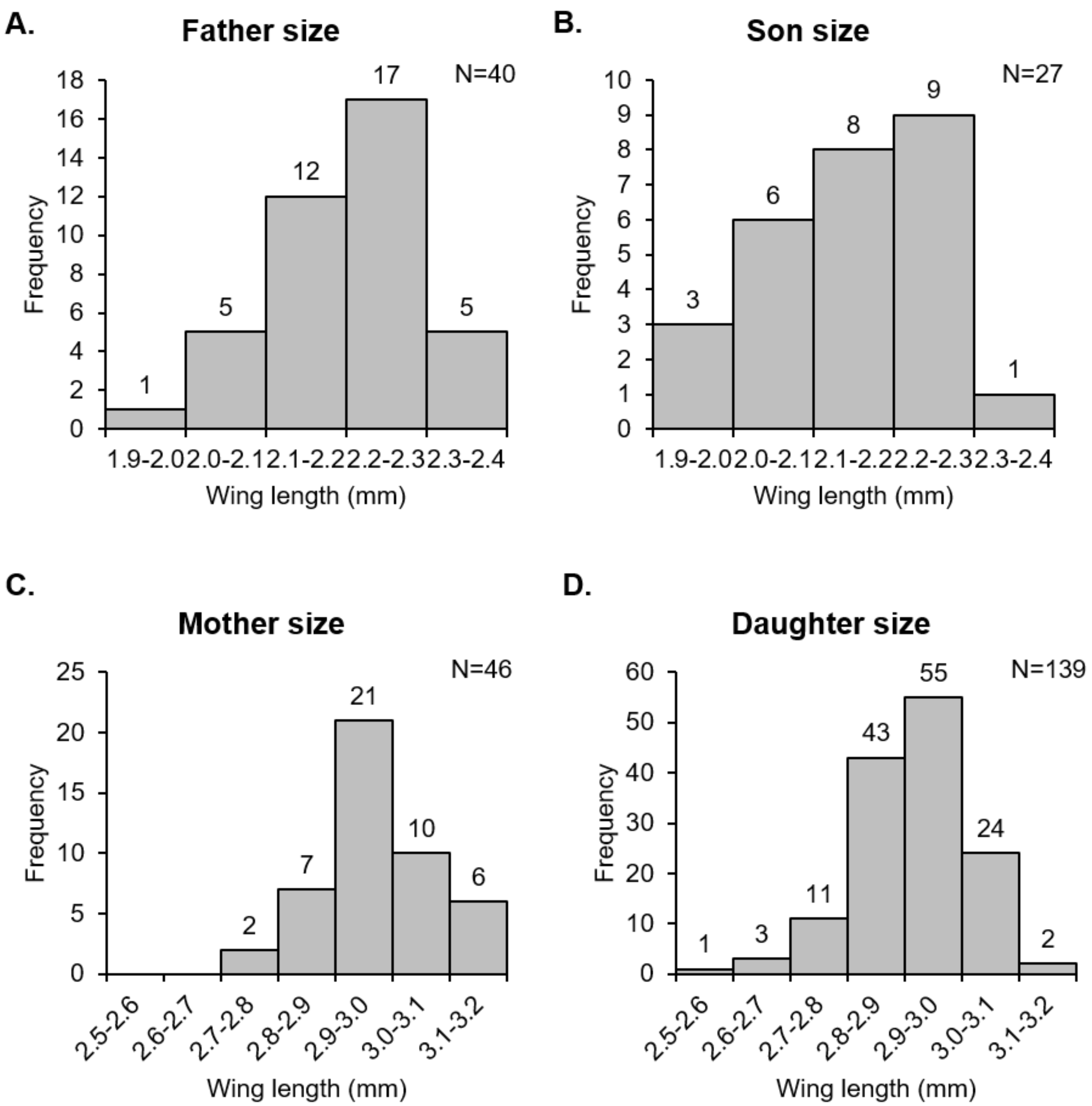


**S4 Figure. Parent and offspring sizes displayed only moderate variation were similar between generations.** Father and son sizes (A, B) were normally distributed and biologically comparable, differing statistically (KW test, P=0.041). Mother, but not daughter size (C, D) was normally distributed and size distributions were similar, despite differing significantly (KW test, P=0.004). Graphs display distribution sample sizes (N) and the number of samples per bin (above bars). For detailed descriptive statistics, see S2 Table.

**S2 Table. Parent and offspring size descriptive statistics reveal only minor variation**. Both parent and offspring size tended to vary only moderately. Normality values display one-sample KS test P-values. Abbreviations: Min, minimum value; Max, maximum value; SD, standard deviation; Var, variance; Sk, skewness.

| Descriptive statistics | | | | | | | | | |
|---|---|---|---|---|---|---|---|---|---|
| | **N** | **Min** | **Max** | **Range** | **Mean** | **Var** | **SD** | **Sk** | **Normality** |
| Father wing length | 40 | 2 | 2.390 | 0.390 | 2.209 | 0.009 | 0.093 | -0.471 | 0.200 |
| Son wing length | 27 | 2 | 2.400 | 0.400 | 2.160 | 0.010 | 0.102 | 0.201 | 0.200 |
| Mother wing length | 46 | 2.770 | 3.190 | 0.420 | 2.982 | 0.010 | 0.100 | 0.085 | 0.200 |
| Daughter wing length | 139 | 2.570 | 3.130 | 0.560 | 2.929 | 0.010 | 0.099 | -0.543 | 0.002 |

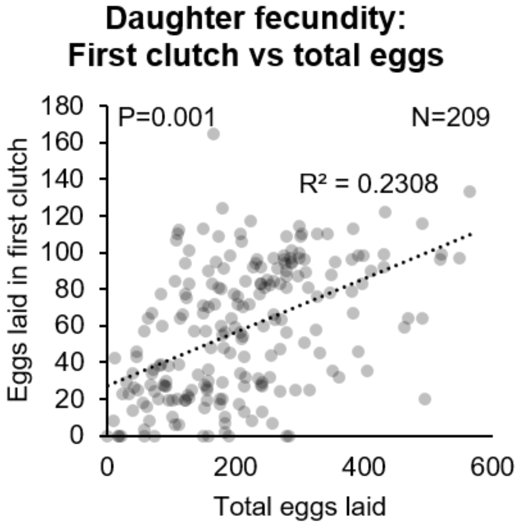


**S5 Figure. Daughter first egg clutch size predicts lifetime fecundity.** Daughters that laid more eggs in their first clutch tended to lay more eggs across their lifetime (P=0.001). Graphs display correlation sample size (N), $R^2$ value for the linear trendline, and LMM P-value.
